# Increased ozone levels disrupt insect sexual communication

**DOI:** 10.1101/2022.08.27.505511

**Authors:** Nan-Ji Jiang, Hetan Chang, Jerrit Weisflog, Franziska Eberl, Daniel Veit, Kerstin Weniger, Bill S. Hansson, Markus Knaden

## Abstract

Insect sexual communication often relies upon sex pheromones^1-3^. Most insect pheromones, however, contain carbon-carbon double bonds and potentially degrade by oxidation^4^. Here, we show that already frequently reported increased levels of Anthropocenic ozone can oxidize all described male-specific pheromones of *Drosophila melanogaster*^5-7^, resulting in reduced amounts of e.g. cis-Vaccenyl Acetate and (*Z*)-7-Tricosene. At the same time female acceptance of ozone-exposed males is significantly delayed. Interestingly, groups of ozonized males also exhibit unnaturally high levels of male-male courtship behavior. When repeating similar experiments with nine other drosophilid species, we observe pheromone degradation and/or corrupted sex recognition in eight of them. Our data suggest that Anthropocenic levels of ozone can oxidize double bonds in a variety of insect pheromones extensively, thereby leading to deviations in sexual recognition.

---

Finding and judging a suitable mate is pivotal for animal reproduction. In this context, most insects use sex pheromones to discriminate conspecifics from allospecifics and to identify the sex and mating status of a potential mate ^1-3^. A particularly well-investigated pheromone is cis-Vaccenyl Acetate (cVA). This compound is produced by male *Drosophila melanogaster*, governs sex recognition, and the cVA amount present on a male has been shown to correlate with the male’s attractiveness to a female ^5,6,8,9^. During copulation, however, the male, in order to ensure its paternity, transfers cVA to the female and thereby reduces the female’s attractiveness to other males ^8,10,11^. cVA is thus attractive to females but repulsive to males. Many fly species of the genus *Drosophila* are known to produce male specific compounds that increase their attractiveness towards conspecific females, become transferred during copulation, and, hence, seem to fulfill similar pheromone-like roles like cVA in *Drosophila melanogaster* ^12,13^. Although being chemically diverse, most of these potential pheromones share one specific feature – they contain carbon double bonds.

During the Anthropocene, insects communicating with such unsaturated pheromones are facing a potential challenge: the oxidization of double bonds by increased levels of oxidant pollutants like ozone ^14^. Pheromone systems have evolved in pre-industrial times with tropospheric ozone values as low as 10 ppb ^15^. However, due to the continuous emission of nitrogen oxides (NOx), volatile organic compounds (VOCs) and the climatic change, the ozone level has already increased to a global yearly average of 40 ppb ^16^. Local extreme ozone levels of over 100 ppb or even 200 ppb have been reported in polluted areas ^17-20^.

Here we show, that already short-term exposure to ozone levels of 100 ppb results in the degradation of many drosophilid pheromones and reduces e.g. the attractiveness of males to females in 7 of 10 tested species. Interestingly, ozone-exposure dramatically increases male-male courtship behavior, probably due to a lack of sex discrimination when male pheromones become degraded.

We first investigated, whether the amount of cVA of *D. melanogaster* males becomes affected by exposure to ozone. Indeed, when comparing with control flies exposed to ambient air (with 4.5±0.5 ppb ozone) only, we found reduced amounts of cVA on flies that were exposed to 100 ppb ozone for 2 hours (Fig. 1, for a schematic of the ozone setup see Fig. S1) and increased amounts of heptanal (Fig. S2), a potential breakdown product of cVA oxidation. Interestingly, many other pheromone compounds like (*Z*)-7-Tricosene (7-T) and (*Z*)-7-Pentasene (7-P)^7,21^, which are known to be involved in reproductive behavior, were decreased after ozone exposure, as well (Fig. 1c).

**Figure 1.**
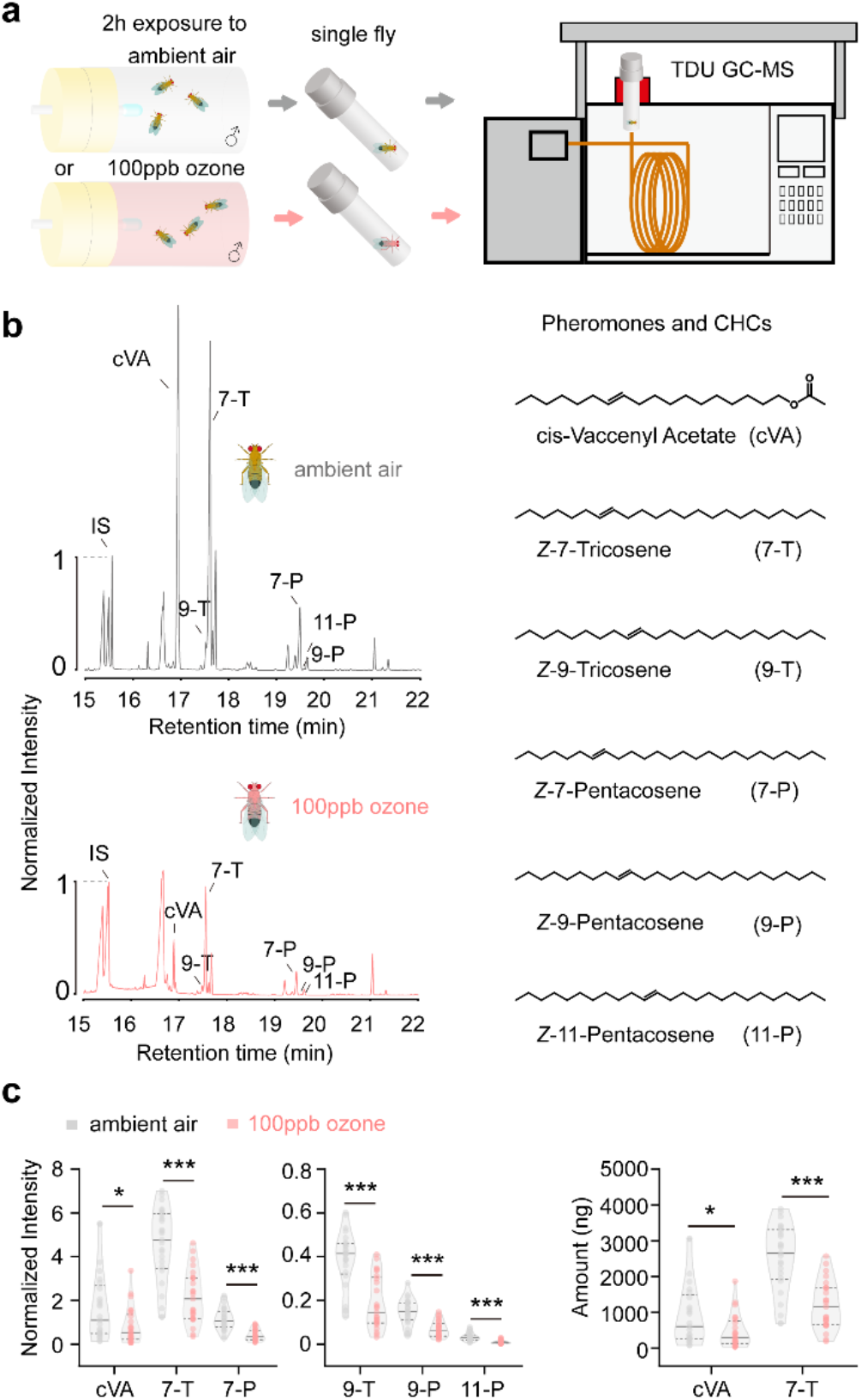
TDU GC-MS analysis of *D. melanogaster* male pheromones and cuticular hydrocarbons. **a**, Schematic drawing of the TDU GC-MS analytical protocol. Male flies were first exposed to ozone or ambient air. Their chemical profile was then analyzed by TDU GC-MS. **b**, Chemical profiles of *D. melanogaster* males after exposure to ozone (pink) or ambient air exposure (grey). Chromatograms (left panel), Chemical structures of male pheromones and curricular hydrocarbons, CHCs (right panel). **c**, Quantitative analysis of male pheromones and CHCs (*t*-test; \**p*<0.05; \*\**p*<0.01; \*\*\**p*<0.001).

We next asked, whether these changes of the males’ chemical profiles would affect their attractiveness to female flies. We therefore exposed male flies either for 30 min to ozone ranging from 50 to 200 ppb or, as a control, to ambient air and afterwards tested their courtship behavior and mating success with non-exposed females in a no-choice mating assay (Fig. 2a).

**Figure 2.**
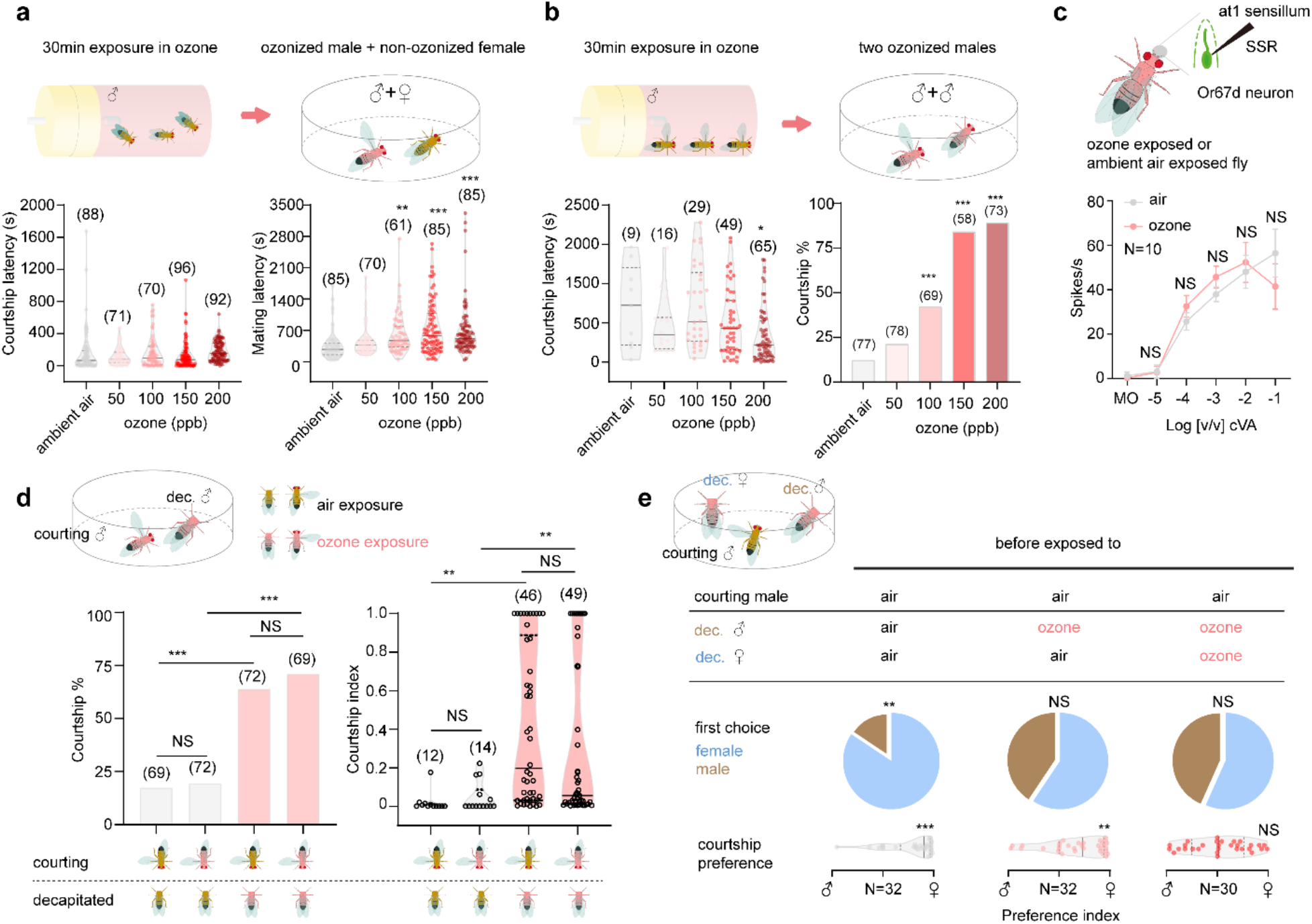
Ozone exposure corrupts *D. melanogaster*’s sex discrimination. **a**, Male/female courtship latency and mating latency after male exposure to different levels of ozone (sample size provided in brackets, *Dunnet’s* test for multiple comparisons against the ambient air control; groups significantly differing from control: \**p*<0.05; \*\**p*<0.01; \*\*\**p*<0.001). **b**, Male/male courtship behavior after both males had been exposed to ozone. Courtship latency, *Dunnet’s* test for multiple comparisons; courtship percentage (percentage of experiments that resulted in courting males), *Fisher’s exact* test with Holm-Bonferroni correction for multiple comparison with control group; significant differences to control depicted as above. **c**, SSR dose response curve of Or67d neuron (at1 sensillum) from male *D. melanogaster*. Top panel, schematic of SSR procedure; data presented as mean ± SE, N=10 for either air or ozone exposure (*t*-test for each concentration; NS indicates no significant difference). Mineral oil (MO) served as the solvent control. **d**, Courtship behavior of an intact *D. melanogaster* male towards a decapitated male. Either the intact, the decapitated or both males were exposed to ozone before the courtship assay. Courtship percentage (see above), *Fisher’s exact* test with Holm-Bonferroni correction for multiple comparison; courtship index (amount of time the intact male courted during the experiment), *Dunnet’s* test for multiple comparisons. **e**, A male fly’s preference for a decapitated male or female. Either the decapitated male, the female or both decapitated flies were exposed to ozone before the courtship assay. First choice presented in pie charts (*Fisher’s exact* test); Preference index (time courting female – time courting male) / total courting time, *t*-test for courtship preference.

Ozone-exposed males did not differ from control males regarding their courtship latency (i.e. the time until they started to court the female; Fig. 2a), and courtship percentage (i.e. the percentage of males that courted females, Fig. S3), showing that male courtship motivation does not seem to be affected by previous exposure to ozone. At the same time, although most males finally mated within the 10min observation (Fig. S3) ozone-exposed males exhibited a longer mating latency than control males (i.e. they needed more time to become accepted by the female; Fig. 2a). Obviously, ozone-exposed males were less attractive to the courted females, which corresponds well with the aforementioned reduction of pheromones upon ozone exposure. We did not observe any effect of ozone when flies were only exposed for 15min, while the mating latency of males again was increased after ozone exposure for two hours or after exposure to higher levels of ozone (150 ppb and 200 ppb) (Fig. S4).

Interestingly, after being exposed to ozone for 2h, males did not recover their original chemical profile and their attractiveness to females after 1 day but exhibited normal pheromone levels and attractiveness after 5 days (Fig. S5).

As mentioned before, male specific pheromones often do not only function as aphrodisiacs for females but also help males to discriminate sexes ^8,9^. We, therefore, hypothesized that ozone exposure of male flies, with the subsequent reduction of their male-specific pheromones, would impede sex discrimination. When we exposed groups of males to 100 ppb of ozone, the results, however, exceeded our expectations. After a brief period (17.72±1.83 min, N=10) of exposure, males started to court each other intensively and to exhibit chaining behavior (supp. Movie S1 and S2), i.e. formed a long chain of courting males that was first described for males carrying a fruitless mutation ^22^. When quantifying such male-male courtship behavior of pairs of ozone-exposed males (i.e. 30 min at 100 ppb) in a no-choice assay (Fig. 2b), we indeed found a higher number of trials that resulted in male courtship as compared to air-exposed control males. Again, the effect of ozone could be increased by either prolonging the exposure time or increasing the level of ozone (Fig. S6). As both males were exposed to ozone in these experiments, we wondered, whether the observed increased male-male courtship was only due to the degraded male-specific pheromones, or was in addition affected by a potential malfunction of sensory neurons, responsible for their detection, due to oxidative stress. Single sensillum recordings (SSR) from antennal trichoid sensilla at1, known to house an olfactory sensory neuron that detects cVA ^9^, revealed similar dose-response curves in ozone exposed flies as in control flies (Fig. 2c). Furthermore, we confronted an intact male with a decapitated male and exposed each of them either to ozone or ambient air before the encounter. We found that whether or not the intact male would exhibit courtship behavior was not influenced by its own exposure to ozone but only by the exposure of the decapitated male (Fig. 2d). Obviously, ozone exposure does not impede the function of pheromone responsive neurons or other levels of signal processing, but rather induces male-male courtship via the degradation of pheromones.

We next asked whether ozone exposure impedes sex-discrimination completely, and analyzed the courting preference of non-ozone exposed males towards a decapitated male and female in a choice assay. The headless flies were exposed to ozone or, as a control, ambient air before. While the males preferentially courted the females in the control experiment, ozone exposure of the decapitated flies resulted in equal courting of female and male flies (Fig. 2e). Interestingly, when only the decapitated male was exposed to ozone before the experiment, the decapitated female was still preferentially courted in the choice assay (Fig. 2e), suggesting that female-specific chemicals are sufficient for sex-discrimination. As the exposure of both sexes to ozone impeded sex-discrimination, also the female cues seem to be sensitive to degradation by ozone. Indeed, when analyzing ozone-exposed females (i.e. 2h at 100 ppb), we found reduced amounts of the described female-specific compounds 7,11-Heptacosadiene (7,11-HD) and 7,11-Nonacosadiene (7,11-ND) ^23,24^ (Fig. S7).

Having shown that exposure to ozone does not only reduce the attractiveness of a *D. melanogaster* male towards females but also strongly compromises sex discrimination, we asked whether this effect is specific to *D. melanogaster* or appears in other species as well. Based on our previous study on male-specific compounds in 99 *Drosophila* species ^13^, we selected 8 further *Drosophila* species that are known to carry male-specific compounds. We exposed their males for 2h to 100ppb ozone and compared their chemical profiles and behavioral performances with that of control males. While seven of these species, except *D. buskii*, exhibited decreased amounts of male-specific compounds after ozone exposure (Fig. 3), most of them showed decreased mating success and/or changes in male-male interactions (Fig. 3). *D. busckii*’s described male-specific compounds do not contain carbon-carbon double bonds and are thus not sensitive to degradation by ozone (Fig. 3). The fact that *D. buskii* males were still less successful in courting females might be due to additional, non-identified but ozone-sensitive compounds governing courtship behavior in this species. Interestingly, both tested *D. mojavensis* subspecies tested, contrary to the other species, exhibited decreased male-male courtship after ozone-exposure. Again additional compounds to those described for these subspecies ^12,13^ might explain these results. We finally tested *D. suzukii*, a species that does not exhibit any sex-specific compounds and whose behavior is supposed to be rather visually driven ^13,25^. As expected, neither mating success nor male-male courtship were affected by ozone treatment (Fig. 3).

**Figure 3.**
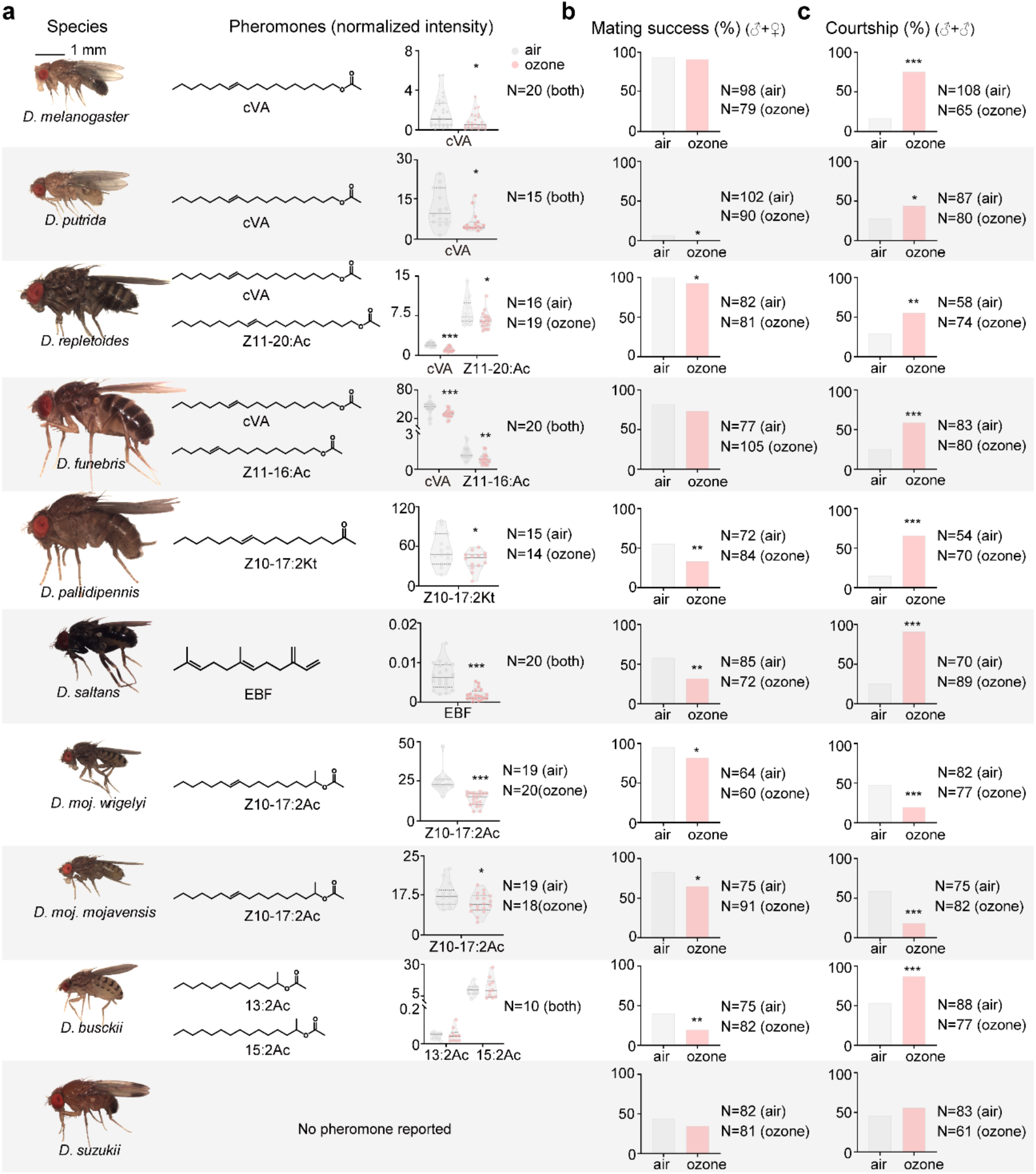
Effects of ozone on male-specific compounds and sexual behavior of 10 drosophilid species. **a**, Data depict normalized peak areas of pheromones in ozone exposed and control flies (*t*-test; \**p*<0.05; \*\**p*<0.01; \*\*\**p*<0.001). **b and c**, Effect on mating success (i.e. percentage of experiments resulting in mating) and effect on male-male courtship behavior (i.e. percentage of experiments resulting in male-male courtship). *Fisher’s* exact test, \**p*<0.05; \*\**p*<0.01; \*\*\**p*<0.001.

## Conclusion

Pollutants like ozone and nitric oxides have been shown to degrade floral volatiles and, hence, corrupt the chemical communication between plants and their pollinating insects ^26-32^. Here we show that ozone in addition can harm insects in another context. The exposure of flies to rather mild ozone levels that have already been reported for polluted areas degrades their unsaturated pheromones and by that affects the sexual communication in 9 out of 10 tested *Drosophila* species. Carbon double bonds, however, are not specific for *Drosophila* pheromones but are a major characteristic of most identified insect pheromones ^33^. As e.g. many female lepidopterans use pheromones for attracting males over long distances, there is ample time for the oxidizing pollutants to potentially degrade the signal before it reaches the receiver. While nowadays the detrimental effect of pesticides on insect populations is well established worldwide ^34,35^, insects obviously face a second problem: the corruption of their chemical information channels by increased levels of oxidizing pollutants.

## Acknowledgments

We thank I. Alali and S. Trautheim for help with the fly breeding.

## Author contributions

NJ.J., B.S.H., and M.K. designed the research plan and N.I.J. performed most of the experiments. H.C. conducted SSR experiments. F.E. and D.V. constructed the ozone exposure device. N.I.J and K.W. analyzed and quantified pheromone compounds. J.W. synthesized synthetic compounds. NJ.J. analyzed experimental data. NJ.J., B. S.H., and M.K. wrote the paper. All authors edited the manuscript.

## Data and materials availability

All data are available in the main text or the supplementary materials.

## Materials and Methods

### Drosophila lines and chemicals stocks

Wild-type flies from the wild-type strain Canton-S used in this study were obtained from the Bloomington 5 Drosophila Stock Center w (NIH P40OD018537). All flies were reared at 25 °C, 12 h Light:12 h Dark and 70% relative humidity. Virgins were collected by using CO_2_ anesthesia. 10-day-old virgin flies were tested in the courtship arena. Care and treatment of all flies complied with all relevant ethical regulations.

### Chemicals

10 We used the described fly compounds including (*Z*)-10-Heptadecen-2-one (Z10-17:2Kt), (*Z*)-11-Hexadecen-1-yl acetate (*Z*11-16:Ac), *rac*-2-Tridecyl acetate (13:2Ac), *rac*-2-Pentadecyl acetate (15:2Ac), (*Z*)-11-Eicosen-1-yl acetate (*Z*11-20:Ac), (*Z*)-10-Heptadecen-2-yl acetate (*Z*10-17:2Ac) by chemical syntheses ^13^. Other chemicals including cis vanccenyl acetate (cVA), (*Z*)-7-Tricosene (7-T), (*Z*)-9-Tricosene (9-T), (*Z*)-7-Pentacosene (7-P), and (*Z*)-9-Pentacosene (9-P), were purchased in high purity from Sigma-Aldrich and Cayman Chemical. (*Z*)-11-15 Pentacosene (11-P) was synthesized via Wittig reaction from bromotetradecane (Sigma-Adrich, Germany) and undecanal (Sigma-Aldrich, Germany). Bromotetradecane (1g, 3.6 mmol) and triphenylphosphine (946 mg, 3.6 mmol) were dissolved in 20 ml of Toluene and heated at reflux for 20 h. After cooling to room temperature, the toluene phase was carefully removed with a pipette. The residue was stirred for 5 min with fresh toluene (10 ml each), before the toluene was pipetted off again. This process was repeated three times with toluene and once with 20 diethylether (10 ml). The highly viscous residue was dried in high vacuum overnight to obtain the Wittig-salt as an amorphous off white solid.

Under argon the Wittig-salt (600 mg, 1.11 mmol) was dissolved/suspended in anhydrous THF (20 ml) and cooled to −30°C. A solution of sodium bis(trimethylsilyl)amide (1.2 ml, 1M in THF) was added dropwise and the mixture was allowed to warm to −10°C. After stirring for 30 min the now deep orange suspension was cooled to −30°C again 25 before adding undecanal (0.23 mL, 1.11 mmol) via syringe. The mixture was stirred overnight at 20°C. The reaction mixture was diluted with n-hexane (20 ml) before adding water (40 ml). The organic phase was removed and the aqueous phase was extracted with n-hexane (2 × 40 ml). The combined organic phases were washed with water and brine (40 ml each) and dried over Mg_2_SO_4_. After filtration the solvent was removed via rotavap, the residue adsorbed onto silica gel and chromatographed with n-hexan/EtOAc (50:1) as eluent to obtain (Z)-11-pentacosene 30 (130 mg, 34% yield based on undecanal) as a colorless oil.

### Ozone exposure system

Ozone was generated by an ozone 100 generator (Aqua Medic, Germany), and then diluted by humidified air to 70% humidity (that before was cleaned from ambient ozone by a Pd scrubber). Concentration of diluted ozone in a 100l 35 Plexiglas container was continuously tracked via a BMT932 ozone monitor (BMT messtechnik GmbH, Germany), and the ozone generator switched on/off automatically to guarantee target concentration in the container. Ozone dilution from the container was blown into the fly tubes containing flies with 95ml/min per tube to expose flies to the different levels of ozone (see Fig. S8).

### TDU GC-MS

To analyze chemical profiles of flies in the TDU GC-MS experiments, 10 days old flies were first exposed to ozone or control air for a given time and afterwards immediately frozen at −20°C for 30min. In order to investigate the recovery of chemical profiles after ozone exposure, males were exposed to 100ppb ozone for 2h, and then moved to tubes with standard food for either 1 or 5 days. Afterwards they were frozen as mentioned before. For TDU GC-MS 45 measurements, individual flies were placed in microvials of thermal desorption tubes (GERSTEL, Germany) and 0.5 µl of C10-Br or C16-Br (10^−3^ dilution in hexane) were added to the microvials as internal standard.

Desorption tubes were transferred using a GERSTEL MPS 2 XL multipurpose sampler into a GERSTEL thermal desorption unit (GERSTEL, Germany). Samples were desorpted at 250 °C for 8 min, then trapped at −50 °C in the liner of a GERSTEL CIS 4 Cooled Injection System, with liquid nitrogen used for cooling. The components were 50 transferred to the GC column by heating the programmable temperature vaporizer injector at 12 °C/s up to 270 °C and then keeping the temperature for 5 min. The GC-MS (Agilent GC 7890A fitted with an MS 5975C inert XL MSD unit; Agilent Technologies, USA) was equipped with an HP5 column (Agilent Technologies, USA). The temperature of the gas chromatograph oven was held at 50 °C for 3 min and then increased by 15 °C /min to 230 °C and then by 20 °C /min to 280 °C, held for 20 min. Mass spectra were taken in EI mode (at 70 eV) in the range from 55 33 m/z to 500 m/z.

### Courtship behavior assays

Virgin males and females of all tested drosophilids were collected after eclosion and raised individually and in groups (20 individuals/vial), respectively. For male-female courtship assays, 15 males were exposed to ozone or ambient air in a vial. Ozone-treated males were then placed with an untreated female into a courtship chamber (see 5 below) and their behavior observed and quantified for 1h. Each courtship arena contained 4 chambers (1 cm diameter × 0.5 cm depth) covered with a plastic slide. Air flow of 0.2 mL/min was added from below to each arena. A GoPro Camera 4 or Logitech C615 was used to record courtship behaviors. Each video was analyzed manually for courtship latency (i.e. the time until the male initiated courtship behavior), courtship percentage (i.e. the percentage of males that showed courtship behavior), courtship index (i.e. the time each male performed courtship behavior 10 during the 10min experimental time), mating latency (i.e. the time until successful mating started), and mating success (i.e. the percentage of males that mated successfully). All behavioral experiments were performed at 25 °C and 70% humidity.

For male-male courtship assays, 25 males were exposed to ozone or ambient air in a fly tube. Afterwards, two males were put into one chamber and their behaviors observed and quantified for 30 min. For *D. melanogaster*, several 15 ozone exposure combinations (i.e.15 min, 30 min, and 2h with 50, 100, 150, 200 ppb, respectively) were tested. Males of other drosophilids were exposed for 2h to 100 ppb. For no-choice assays with decapitated males, males were exposed to ozone, and then decapitated. An intact male and a decapitated male were put into one chamber and the courtship behavior of the intact male observed and quantified for 30 min. For the two-choice assays with decapitated flies, males and/or females were exposed to ozone, and then decapitated. An intact male, and a 20 decapitated male and female were put into one chamber. The preference index of the courting intact male was calculated as (time of intact male court to decapitate female - time of intact male court to decapitate male)/30 min.

### Single sensillum recording (SSR)

Male *D. melanogaster* flies were exposed to either ozone or ambient air, and then immobilized in a pipette tip. A 25 reference electrode was put into the eye; another tungsten electrode was inserted into the target sensillum. The at1 was identified based on their location and spontaneous activities. Signals were amplified by Syntech Universal AC/DC Probe (Syntech, Germany), sampled (96,000.0 samples/s), and filtered (500-5000 Hz with 50/ 60 Hz suppression) via a USB-IDAC (Syntech, Germany) connection to a computer. Action potentials were extracted using AutoSpike software (Syntech, Germany). Synthetic compounds were diluted in mineral oil (MO) (Sigma-30 Aldrich, Germany). Before the test, 10 µl of the diluted odor was freshly loaded onto a small piece of filter paper (1 cm^2^), and placed inside a glass Pasteur pipette. The tested odor dosages were ranging from 10^−5^-10^−1^ dilution (v/v). The odorant was delivered by inserting the tip of the pipette into a constant, humidified airstream flowing at 600 ml/min through a stainless steel tube (diameter, 8 mm) ending in 1 cm distance from the antenna. Neural activity was recorded for 10 s, starting 3 s before the stimulation period of 0.5s. Responses from individual neurons were 35 calculated as the increase (or decrease) in the action potential frequency (spikes/s) relative to the pre-stimulus frequency. Traces were processed by sorting spike amplitudes in AutoSpike, analysis in Excel and illustration in Adobe Illustrator CS (Adobe systems, USA).

### Statistical analyses

40 Statistical analyses (see the corresponding legends of each figure) and preliminary figures were conducted using GraphPad Prism v. 8 (GraphPad Software, USA). Figures were then processed with Adobe Illustrator CS5.

## Supplementary Materials for

**Fig. S1.**
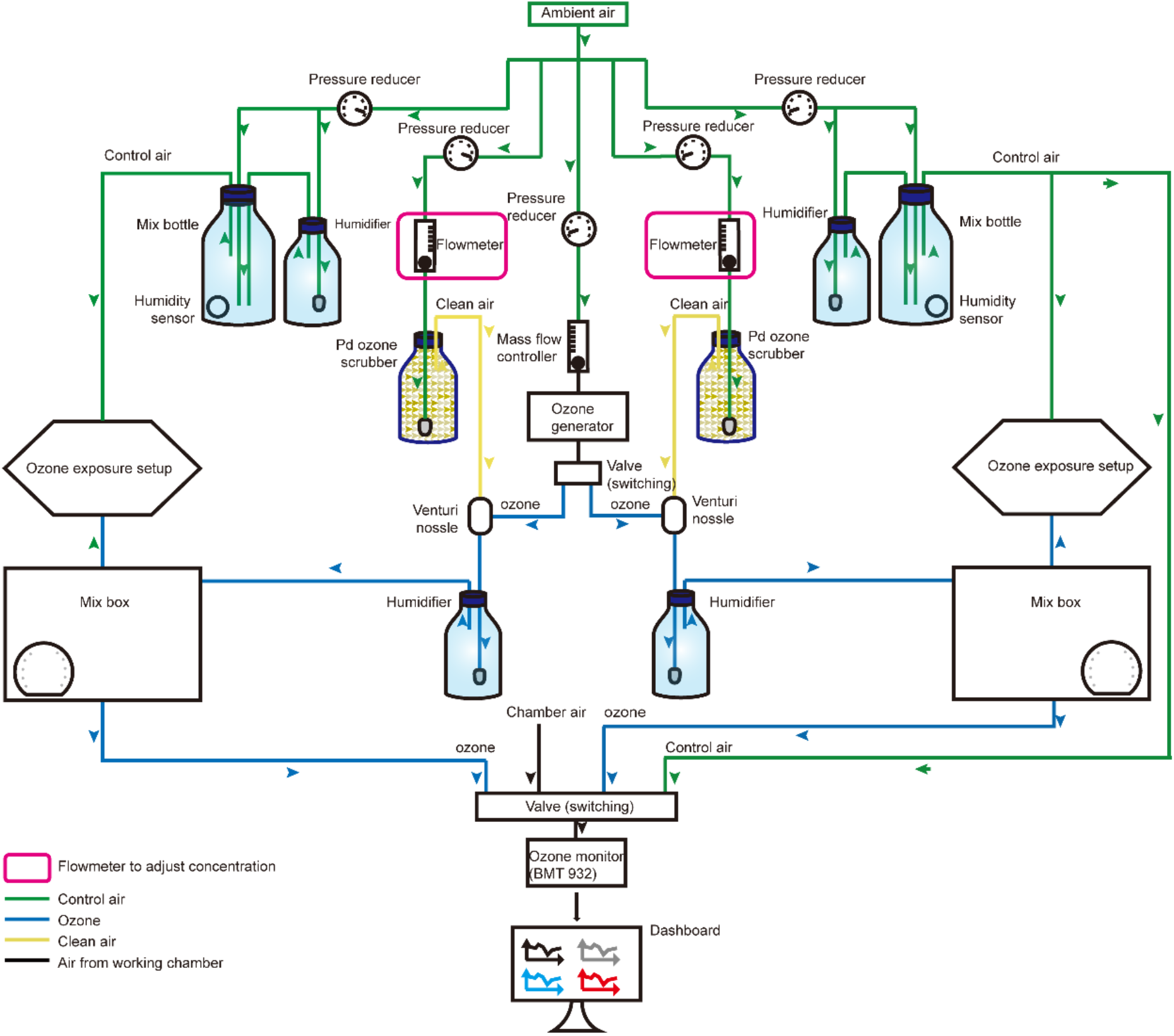
The ozone exposure system. Schematic of the device to produce defined levels of ozone concentrations.

**Fig. S2.**
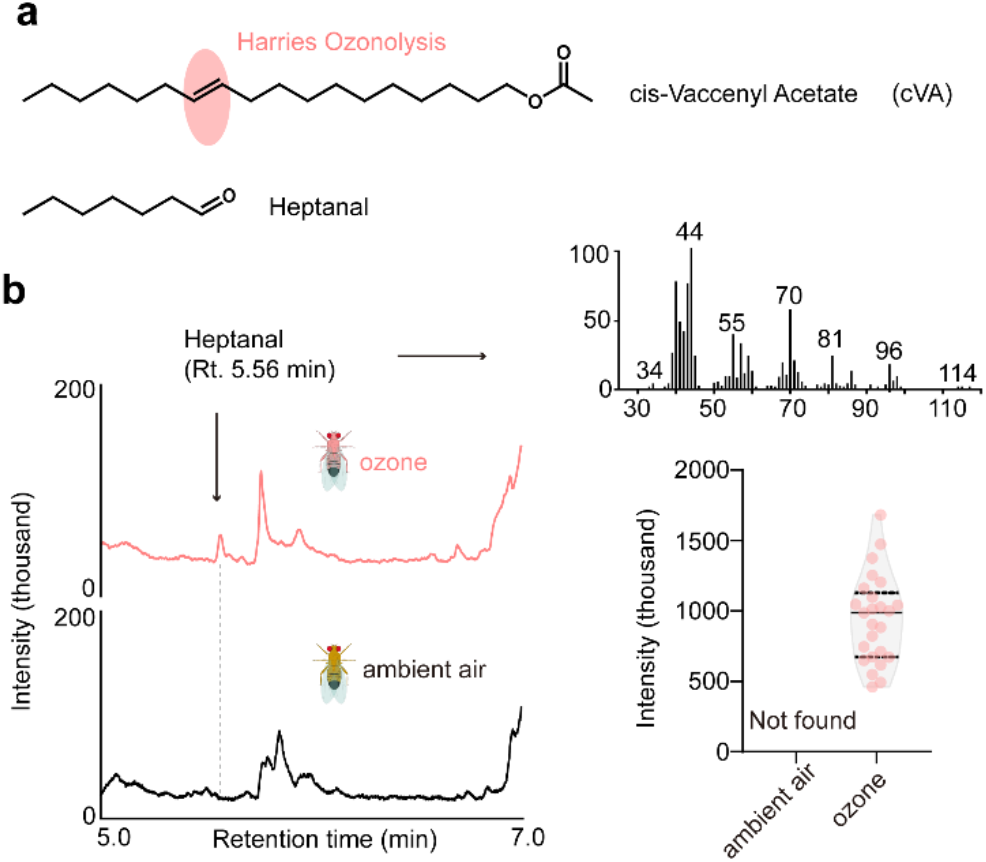
Ozone-exposed males emit more heptanal. **a**, Potential degradation of cVA following Harries Ozonolysis. **b**, Chromatograms (left) and MS spectrum (right up) and quantification (right down) of heptanal from males exposed to ozone or ambient air.

**Fig. S3.**
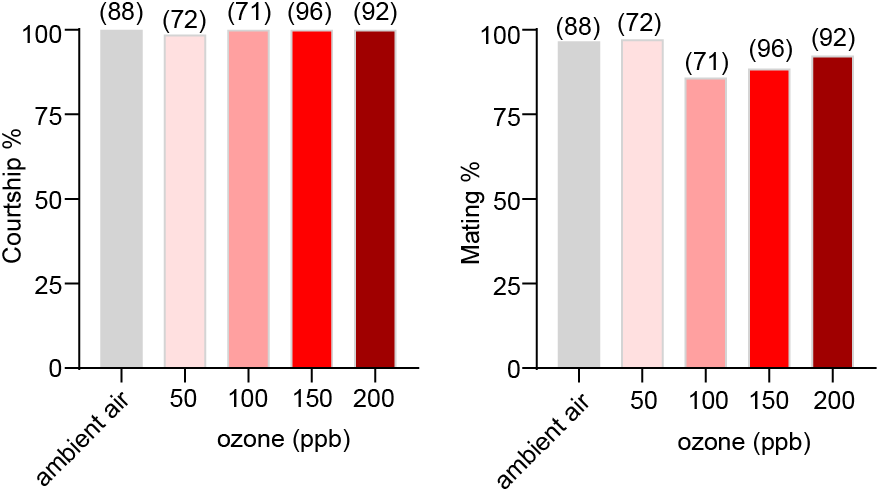
Effect of male exposure for 30 min to different levels ozone on courtship behavior. Courtship percentage (i.e. percentage of experiments resulting in courtship behavior) and mating success (i.e. percentage of experiments resulting in mating). Sample sizes provided in brackets, no significant differences from control ambient air control group (*Fisher’s exact* test with Holm-Bonferroni correction for multiple comparison with control group).

**Fig. S4.**
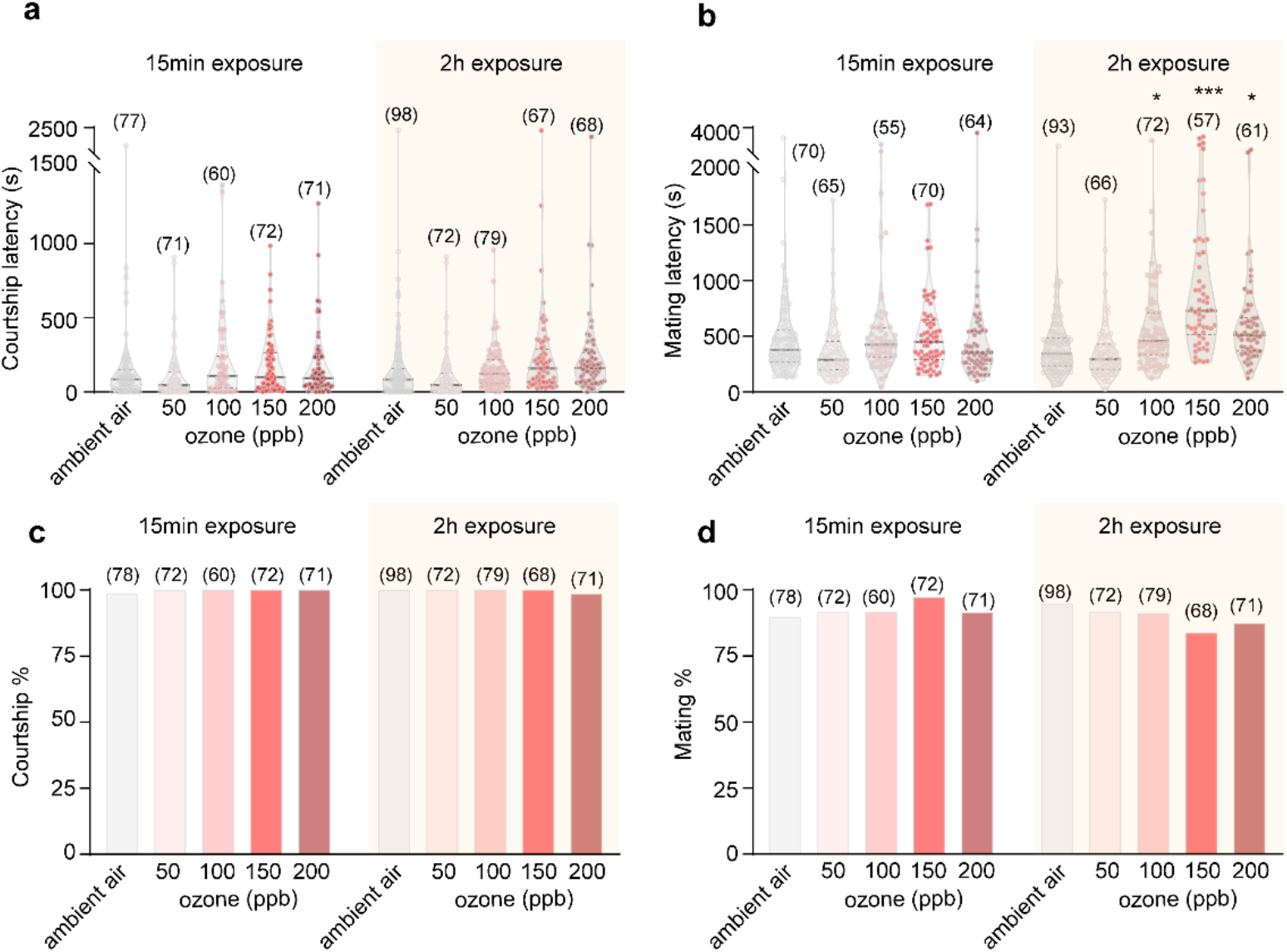
Effect of exposure time and level of ozone on courtship behavior. **a**, Courtship latency (i.e. time until male starts courting). **b**, Mating latency (i.e. time until mating starts). **c**, Courtship percentage (i.e. percentage of experiments resulting in courtship behavior). **d**, Mating success (i.e. percentage of experiments resulting in mating). Samples are given in brackets. (**a** and **c**, *Dunnet’s* test for multiple comparisons against ambient air control group; **b** and **d**, *Fisher’s exact* test with Holm-Bonferroni correction for multiple comparison with control group, \**p*<0.05; \*\**p*<0.01; \*\*\**p*<0.001).

**Fig. S5.**
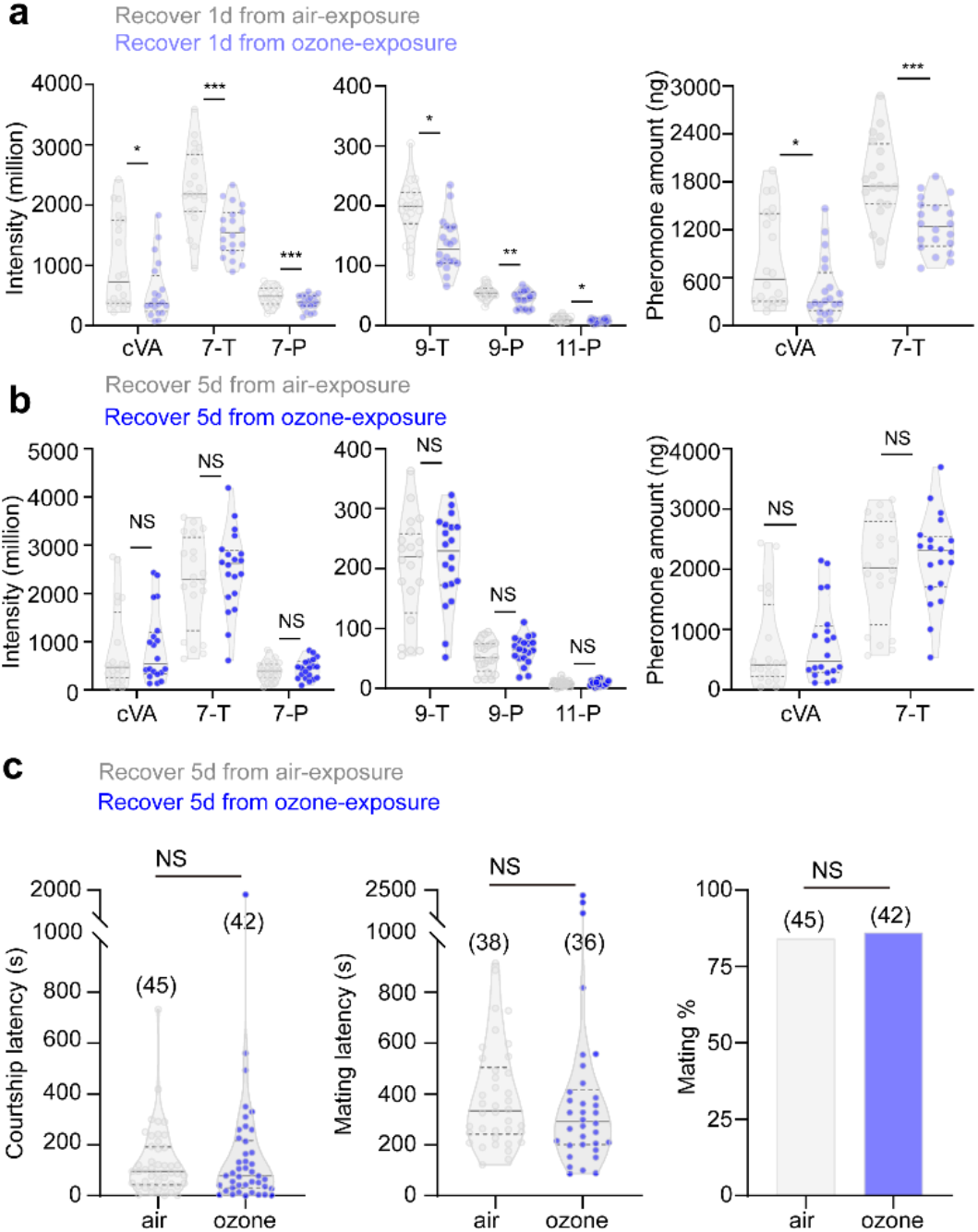
Recovery of chemical profiles after ozone exposure. **a**, One day after exposure, male compounds of ozone-exposed flies still differ from control males. **b**, Five days after exposure, chemical profiles are fully 5 recovered. cVA: cis-vaccenyl acetate; 7-T: (*Z*)-7-Tricosene; 9-T: (*Z*)-9-Tricosene; 7-P: (*Z*)-7-Pentacosene; 9-P: (*Z*)-9-Pentacosene; 11-P: (*Z*)-11-Pentacosene. Data depict peak area for the different described pheromones of *D. melanogaster* males and the quantified amount of cVA and 7-T. **c**, Five days after exposure courtship behavior is fully recovered. (**a-c**) *t*-test; NS indicates no significant difference, \**p*<0.05; \*\**p*<0.01; \*\*\**p*<0.001.

**Fig. S6.**
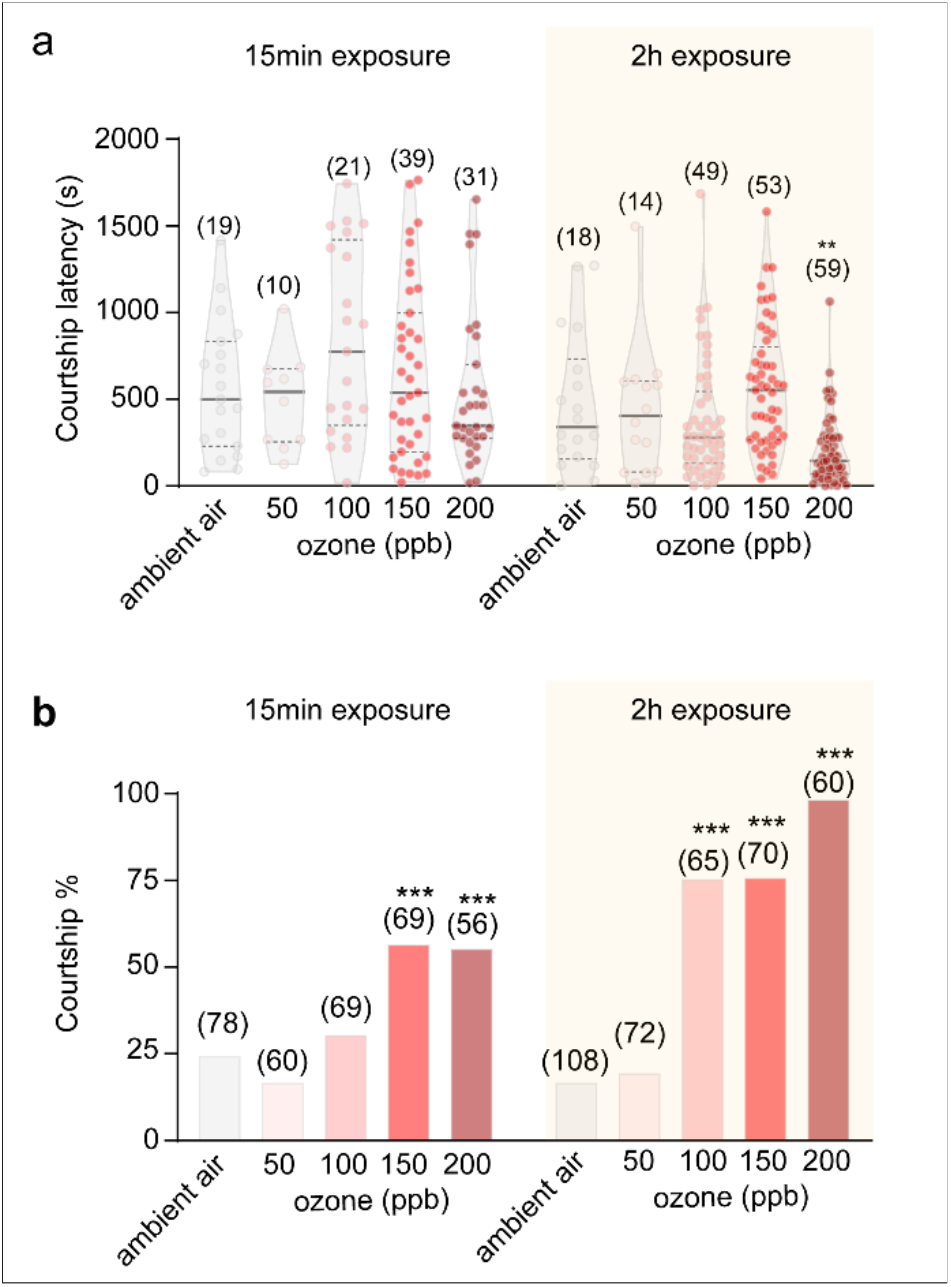
Ozone induces *D. melanogaster* male-male courtship behavior. **a**, Courtship latency (i.e. time until courtship behavior starts) after 15min (left panel) or 2h (right panel) exposure to different levels of ozone. Sample sizes are provided in brackets. *Dunnet’s* test for multiple comparisons against the ambient air control; group significantly differing from control: \*\**p*<0.01. **b**, Courtship percentage (i.e. percentage of experiments that result in courtship within 10min). *Fisher’s exact* test with Holm-Bonferroni correction for multiple comparison with control group, \*\*\**p*<0.001.

**Fig. S7.**
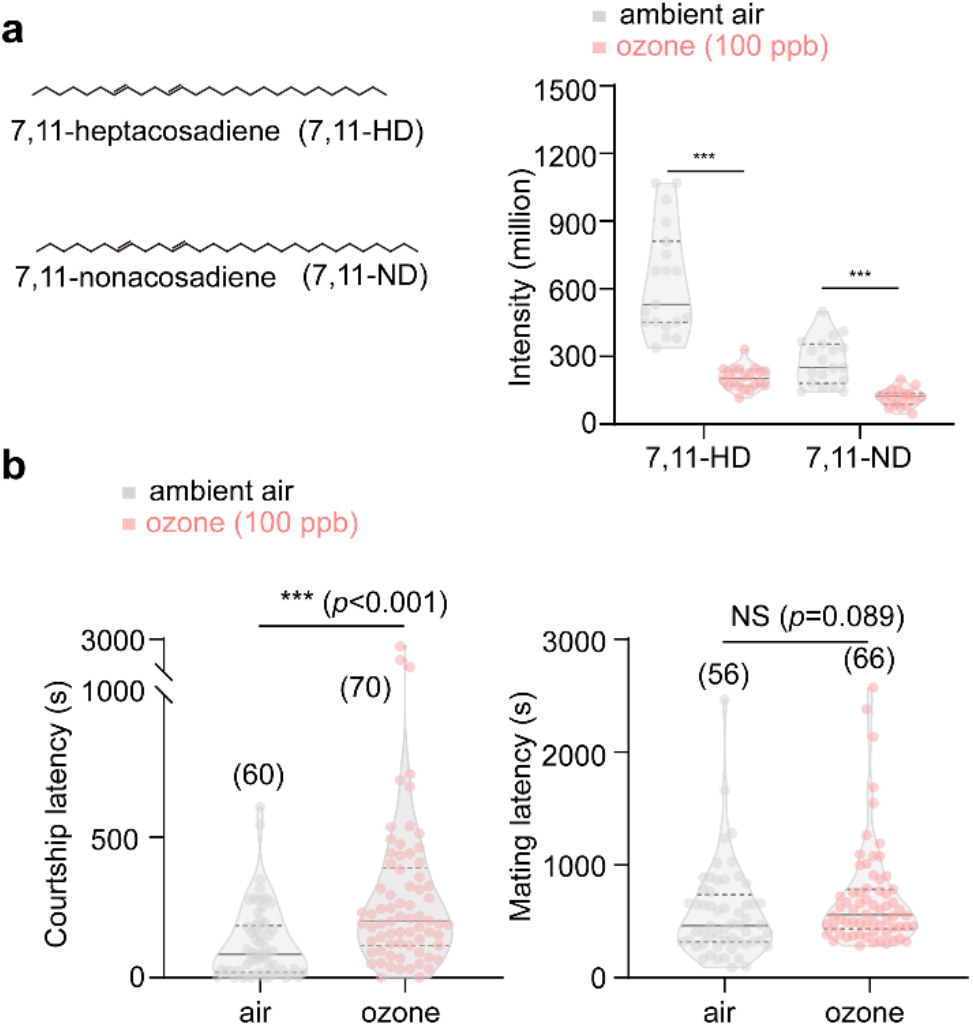
Exposure of female flies to ozone changes their chemical profiles and attractiveness to males. **a**, Peak area of 7,11-Heptacosadiene (7,11-HD) and 7,11-Nonacosadiene (7,11-ND) in ozone exposed and control flies (*t*-test; \*\*\**p*<0.001). **b**, Courtship and mating latency of males confronted with ozone-exposed and control females (*t*-test; NS indicates no significant difference; \*\*\**p*<0.001).

**Movie S1**.

Courtship-chain formation of *D. melanogaster* males during ozone exposure. Males were exposed to 100 ppb ozone for ca. 20 min.

**Movie S2**.

No male-male courtship of *D. melanogaster* males during exposure to control air.

